# Condensate-Driven Transcriptional Reprogramming Defines Core Vulnerabilities in Esophageal and Gastric Cancers

**DOI:** 10.64898/2026.02.23.707358

**Authors:** Luis Alvarez-Carrion, Andrés R. Tejedor, Juan A. Ardura, Verónica Alonso, Carlos Alonso-Moreno, Rosana Collepardo-Guevara, Irene Gutierrez-Rojas, Cristian Privat, Víctor Moreno, Emiliano Calvo, Balazs Gyorffy, Jorge R. Espinosa, Alberto Ocana

**Affiliations:** Cátedra INTHEOS-START-CEU de Oncología de precisión, Facultad de Medicina, Universidad San Pablo-CEU, Montepríncipe, 28660 Boadilla del Monte, Madrid, Spain; Department of Physical Chemistry, Universidad Complutense de Madrid, Av. Complutense s/n, Madrid 28040, Spain; Instituto Pluridisciplinar, Universidad Complutense de Madrid, P.° de Juan XXIII, 1, Moncloa - Aravaca, 28040 Madrid, Spain; Yusuf Hamied Department of Chemistry, University of Cambridge, Lensfield Road, Cambridge CB2 1EW, UK; Universidad de Castilla-La Mancha, Departamento de Química Inorgánica, orgánica y bioquímica. Facultad de Farmacia-Centro de Innovación en Química Avanzada (ORFEO-CINQA), Unidad nanoDrug, 02008 Albacete, Spain; Department of Genetics, University of Cambridge, Cambridge, UK; Experimental Therapeutics CRIS Cancer Unit, Hospital Clinico San Carlos and IdSCC and CIBERONC, Madrid, Spain; START Madrid-Fundación Jiménez Díaz (FJD) Early Phase Program, Fundación Jiménez Díaz Hospital, Madrid, Spain; START Madrid-CIOCC, Centro Integral Oncológico Clara Campal, 28050 Madrid, Spain; Dept. of Bioinformatics, Semmelweis University, H-1094, Budapest, Hungary; Institute of Transdisciplinary Discoveries, Medical School, University of Pecs, H-7624, Pecs, Hungary; Cancer Biomarker Research Group, Institute of Molecular Life Sciences, HUN-REN Research Centre for Natural Sciences, H-1117, Budapest, Hungary; PhAsIca Biosciences S.L, Calle Velázquez, 27, 28001 Madrid, Spain

**Author notes:** JRE and AO are both corresponding authors. LA and ART are both co-first authors.

## Abstract

Biomolecular condensates organize key nuclear functions by compartmentalizing biomolecules, yet their contribution to gastrointestinal tumorigenesis remains poorly defined. Integrating multi-omics profiling, functional genomics, and molecular dynamics simulations, we reveal that esophageal and gastric cancers share a condensate-enriched transcriptional program driven by intrinsically disordered proteins involved in transcription, RNA processing, and replication stress. Transcriptomic analyses identify a hyperactive transcriptional state with upregulation of condensate-associated genes, including *TOPBP1* and *CHERP*. Dependency mapping demonstrates that these proteins are essential for tumor cell viability, defining a conserved condensate core across different tumor types. Machine-learned predictions and residue-resolution coarse-grained simulations confirm that *TOPBP1* and *CHERP* undergo phase separation through homotypic interactions mediated by intrinsically disordered regions, with saturation concentrations below 2 µM, consistent with spontaneous condensate formation observed *in vitro.* Together, these findings establish condensate organization as a fundamental mesoscale principle in upper gastrointestinal cancers and nominate condensate scaffolds as tractable therapeutic vulnerabilities.

## Introduction

Cells are organized not only by membrane-bound organelles but also by biomolecular condensates—dynamic, membraneless compartments formed through phase separation ^1,2^. These assemblies compartmentalize biochemical reactions, concentrate macromolecules, and enable rapid responses to environmental cues ^3–5^. Their formation relies on multivalent, transient interactions among proteins and nucleic acids, frequently mediated by intrinsically disordered regions (IDRs) or low-complexity domains (LCDs) ^6–9^. Through this biophysical principle, condensates regulate diverse cellular processes, including transcription, RNA metabolism, DNA repair, and signal transduction, conferring a higher order dynamic functional organization to the cell^10–12^.

Aberrant condensate formation has recently been implicated in cancer ^13–16^. By reorganizing chromatin architecture, transcriptional machinery, or stress-response pathways, condensates can sustain oncogenic signaling, protect cancer cells from metabolic stress, and promote drug resistance ^10,17^. Consistent with this view, recent studies have begun to directly link condensate dysregulation to tumor initiation, progression, and therapeutic response across multiple cancer contexts ^17–22^. These emergent properties redefine the concept of druggability, suggesting that modulation of condensate composition or material properties could offer a new therapeutic frontier in precision oncology ^23^. Despite these advances, how condensate biology contributes to specific cancer types remains largely unexplored.

Within the cancer spectrum, esophageal and gastric cancers constitute a class of exceptionally aggressive malignancies, characterized by deep molecular heterogeneity and unfavorable clinical outcomes ^24,25^. Although genomic studies have identified multiple genomic alterations in esophageal and gastric cancers, the mechanistic principles that sustain tumor robustness and therapeutic resistance remain poorly understood, underscoring the need for integrative approaches that link genetic, epigenetic, and biophysical alterations to functional tumor behavior ^17–22,26,27^. Although recent studies have tentatively explored the therapeutic relevance of biomolecular condensates in gastric and esophageal cancers ^28–31^, these efforts merely scratch the surface, highlighting the absence of a systematic, computational and biophysically informed framework to dissect condensate-driven vulnerabilities in these diseases ^29–31^.

Addressing this knowledge gap requires synergic experimental and computational approaches capable of capturing both the physical principles and functional consequences of biomolecular condensate formation and disolution ^32^. Multiple experimental techniques have been widely applied to condensates to provide direct insight into phase-separation propensity, condensate material properties, and dynamic regulation, enabling quantitative assessment of condensate nucleation, stability, and responsiveness to environmental or molecular perturbations ^33–37^. In particular, *in vitro* reconstitution assays offer a powerful and well-controlled framework to unravel condensate formation, allowing systematic variation of amino acid sequence, protein concentration, and physicochemical conditions ^38–41^. These approaches can be complemented by advanced imaging and spectroscopic techniques that access the relevant length and time scales of phase separation and coalescence, including super-resolution microscopy such as stimulated emission depletion (STED) ^42,43^, fluorescence correlation spectroscopy (FCS) ^40,44^, fluorescence recovery after photobleaching (FRAP) ^45–47^, or raster image correlation spectroscopy (RICS)^48,49^, which together enable quantitative characterization of condensate formation, internal mobility, and material properties.

Complementing experimental approaches with machine-learned predictors and molecular dynamics (MD) simulations offer a mechanistic framework to interpret experimental observations and to explore regimes that are difficult to access experimentally ^50–53^. In particular, atomistic simulations resolve sequence-specific interactions and conformational landscapes^54,55^, whereas coarse-grained models extend accessible length and time scales, allowing quantitative mapping of phase diagrams, saturation concentrations, and emergent condensate material properties ^51,56,57^. Together, these complementary approaches lay the foundation for AI-driven frameworks that integrate molecular, functional and biophysical data to uncover condensate-driven cancer dependencies.

In this study, we present an AI-driven discovery platform that integrates multi-omics and biophysical data to identify condensate-associated oncogenic dependencies. The platform combines large-scale transcriptomic profiling, functional dependency datasets, and residue-level chemically-specific computational modeling of phase-separation propensity, enabling the prioritization of genes that are simultaneously upregulated, essential for tumor cell viability, and intrinsically prone to condensate formation. This integrative framework bridges molecular expression patterns to mesoscale condensate organization and facilitates the discovery of new therapeutic targets rooted in condensate dynamics.

Using this approach, we identify a set of genes recurrently overexpressed in esophageal and gastric cancers, essential for tumor maintenance, and validated as contributors to condensate formation. Integration of machine-learning algorithms and residue-level coarse-grained MD simulations identifies the proteins *TOPBP1,* and *CHERP* as oncogenic drivers that form biomolecular condensates. Together, these findings establish biomolecular condensates as a central organizational in upper gastrointestinal malignancies and position our platform as a powerful tool to uncover condensate–driven vulnerabilities for future therapeutic exploitation.

## Material and methods

### Data sources and inclusion criteria

RNA-sequencing (RNA-seq) read count data were obtained from two public repositories. Normal tissue controls were retrieved from the Genotype-Tissue Expression (GTEx) portal. Tumor and related samples were downloaded from the GDC portal ^58^(TCGA projects, HTSeq-Counts files). Only primary tumors and normal were retained for analysis; non-primary and non-human samples were excluded.

### Preprocessing and normalization

HTSeq-Counts/ read count files were processed with the DESeq2 normalization pipeline (DESeq2 algorithm for within-sample size factor normalization)^59^, followed by an additional global scaling normalization to render expression levels comparable across cohorts. After normalization, per-gene mean and median expression values in tumor tissues were computed. A tumor-expression filter requiring both mean and median >1,000 (counts) was applied in specific downstream lists to restrict attention to genes with substantial absolute expression (cut-off chosen as the approximate per-sample mean expression across all genes).

### Statistical analysis

Differential expression between normal and tumor samples for each histology was assessed by the nonparametric Mann–Whitney U test (two-sided). Genes were ranked by fold change (tumor/normal) for reporting. The analysis was performed for esophageal carcinoma with n_normal_ = 418 and n_tumor_ = 161 and stomach adenocarcinoma with n_normal_ = 294 and n_tumor_ = 375. For each tumor all genes were saved in a ranked order by fold change together with Mann–Whitney p-value, mean and median expression in normal and tumor groups.

### Identification of biomolecular condensate–related genes

To define the set of genes associated with biomolecular condensate biology, we selected Gene Ontology (GO) terms encompassing biological processes and subcellular compartments previously linked to condensate formation (Date Oct 1^st^, 2025). The selected GO categories included *positive regulation of transcription by RNA polymerase II (GO:0045944)*, *regulation of DNA-templated transcription (GO:0006355)*, *chromatin remodeling (GO:0006338)*, *chromatin organization (GO:0006325)*, *transcription by RNA polymerase II (GO:0006366)*, *RNA processing (GO:0006396)*, and *RNA splicing (GO:0008380)*. In addition, condensate-associated organelles and regulatory assemblies were incorporated, including *nuclear speck (GO:0016607)*, *P-body (GO:0000932)*, *PML body (GO:0016605)*, *Cajal body (GO:0015030)*, *cytoplasmic stress granules (GO:0010494)*, and *centrosome (GO:0005813)*. Terms related to protein quality control and signaling, such as *ubiquitin-dependent protein catabolic process (GO:0006511)*, *proteolysis involved in cellular protein catabolic process (GO:0051603)*, *intracellular signal transduction (GO:0035556)*, and *signal transduction (GO:0007165)*, were also included. The union of genes annotated under these GO terms constituted the condensate-related gene set used for subsequent transcriptomic and functional analyses

### Dependency analysis using DepMap

The essentiality of candidate targets in esophageal and gastric cancer cell lines was evaluated using the DepMap portal (https://depmap.org/portal/; accessed November 2025). This platform integrates large-scale functional genomics datasets to assess gene dependency profiles derived from CRISPR-Cas9 (CERES scores)^60^ and RNA interference (DEMETER2 scores)^60^ screens. Genes were categorized as *common essential* or *strongly selective* according to their dependency patterns across cell lineages. A mean dependency score threshold of –0.7 was applied to identify genes whose inhibition approaches cellular essentiality.

### Gene expression analysis using TCGA

The expression of CHERP and TOPBP1 was analyzed across all tumor types using data from The Cancer Genome Atlas (TCGA) accessed via cBioPortal (https://www.cbioportal.org/; accessed November 2025). Normalized RNA-sequencing expression values, reported as transcripts per million (TPM), were retrieved and compared between tumor and matched normal tissues. Differential expression was calculated using log_2_(TPM + 1)–transformed values, and genes showing consistent upregulation in tumors were prioritized for further analysis. For downstream evaluations, a mean TPM value ≥ 32 was considered indicative of medium-to-high expression, corresponding to biologically relevant transcriptional activity in malignant tissues.

### Predicting phase separation propensity and molecular dynamics validation

To identify protein sequences with intrinsic capacity of forming biomolecular condensates via homotypic phase separation, we employed a Machine Learning (ML) predictor^61^ trained on coarse-grained MD simulations using the CALVADOS2 force field^62^. The model quantitatively estimates the transfer free energy associated with partitioning between the condensed and dilute phases, and the corresponding saturation concentration (C_sat_) for single-component condensates at 293 K and 150 mM of NaCl directly from protein sequence. Importantly, this predictor was developed and validated specifically for IDRs. Accordingly, for the purpose of the initial C_sat_ prediction analysis, we restricted the input sequences to IDRs within each protein of interest. IDRs were identified using AlphaFold3 (AF3) ^63^ confidence scores. Residues with predicted local distance difference test (pLDDT) scores below 70 were classified as disordered, and proteins were required to contain at least 40% of disordered regions to be included in the analysis. For each selected protein, only the amino acids belonging to the AF3-predicted IDRs were used as input to the ML model, yielding an estimated C_sat_ dictated exclusively by IDR–IDR homotypic interactions. As the predictor does not account for folded domains, the C_sat_ values capture only the intrinsic contributions of IDRs to phase separation. In full-length proteins, folded domains may either: (1) attenuate phase separation by intramolecularly sequestering IDRs, thereby rendering the IDR-based estimate a lower bound on C_sat_; or (2) enhance phase separation by contributing additional multivalent interactions, in which case the IDR-based estimate represents an upper bound.

To extend the analysis beyond isolated IDRs and capture folded domain contributions, we performed Direct Coexistence (DC) MD simulations of full-length *CHERP, TOPBP1, HCFC1*, and *IRS2* using the Mpipi-Recharged model^64^ in LAMMPS^65^. Mpipi-Recharged is a physics-based implicit-solvent, residue-level coarse-grained model for protein and RNA condensates. The model has demonstrated near-quantitative agreement with experimental phase diagrams and has been successfully applied to intrinsically disordered, multidomain, and highly charged protein systems^64,66–68^. The model enables simulations of full-length proteins by representing folded domains as rigid bodies covalently linked to flexible disordered segments. Previous benchmarking for IDRs has shown that Mpipi-Recharged and CALVADOS 2 yield consistent predictions of phase behavior for disordered proteins, supporting the complementarity of these approaches^62,66,68^.

In Mpipi-Recharged, each amino acid or nucleotide is represented by a single bead connected by harmonic bonds to residues adjacent in the sequence. To describe multi-domain proteins, residues with pLDDT > 90 were classified as globular and modeled using rigid bodies, with beads positioned at the corresponding Cα coordinates from AlphaFold (AF v6) structures (UniProt IDs: Q8IWX8 for *CHERP*, Q92547 for *TOPBP1*, P51610 for *HCFC1*, and Q9Y4H2 for *IRS2*). IDRs were modeled as fully flexible polymers covalently linked to the rigid domains. Non-bonded interactions consist of the sum of the hydrophobic and electrostatic interactions. Hydrophobic interactions are described by the Wang–Frenkel (WF) potential, which describes short-ranged excluded-volume repulsion and mid-ranged attraction^69^. To account for buried residues within folded domains, hydrophobic interactions involving globular regions were reduced: WF interactions between flexible and globular residues were scaled by √0.7 and globular–globular by a factor of 0.7. Electrostatic interactions were modeled using a Yukawa potential, allowing pair-specific modulation of screened Coulomb interactions. (see ref.^64^ for full force field details).

Using this model and the LAMMPS package, we run DC simulations at 280K and 150 mM to elucidate the phase behaviour of *CHERP, TOPBP1, HCFC1*, and *IRS2*. We placed several replicas of the same protein in an orthogonal slab box with periodic boundary conditions. Given the length of the proteins (some above 2000 amino acids), we use a total of 25 protein replicas in each simulation box. The dimensions of the short sides of the slab box were set to be larger than twice the mean radius of gyration of each protein, specifically we use a section of 200×200 Å for CHERP and IRS2, 225×225 Å for TOPBP1, and 240×240 Å for HCFC1. Simulations are carried out in the canonical (NVT) ensemble using a Nosè-Hoover thermostat for the rigid bodies (representing the globular domains) included in the RIGID package, and a Langevin thermostat for the particles in flexible regions, both with a relaxation time of 5 ps. The time step for the Verlet integration of the equations of motion is 10 fs. After an equilibration period (∼10ns), we run the simulation for 2 *μ*s depending on the specific system. Critical solution temperatures are calculated using the law of rectilinear diameters and critical exponents ^66^:

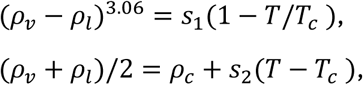

where *ρ*_*v*_ and *ρ_l_* are the densities from the diluted and condensed phases, respectively, 𝑇_𝑐_ is the solution critical temperature, *ρ*_𝑐_ is the critical density, and 𝑠_1_ and 𝑠_2_ are fitting parameters.

## Results

### RNA-Seq Analysis Identifies Shared Transcriptional Deregulated Genes in Esophageal and Gastric Cancers

To detect potential genes that code for proteins that form biomolecular condensates we first performed a comprehensive RNA-seq analyses of primary esophageal and gastric tumors searching for upregulated transcripts. This study revealed strikingly similar distributions of differential gene expression across both malignancies, encompassing 21,479 genes. In esophageal tumors, 19% of these genes (4,093 genes) exhibited high expression levels (median >1,000 TPM), and 4.6% (992 genes) were significantly deregulated relative to matched normal tissues with a clear predominance of upregulated (4.1%) over downregulated (0.5%) transcripts. (Fig. 1 A, B and C). Gastric tumors mirrored this trend, displaying 17.7% (3,803 genes) with high expression and 4.3% (919 genes) showing significant transcriptional deregulation, again dominated by upregulated genes (3.8% vs. 0.4%). Only a limited fraction (<1%) of genes in either tumor type showed strong transcriptional shifts (fold change > 4), consistent with a widespread but quantitatively moderate reprogramming of the tumor transcriptome (Fig. 1 D, E and F).

**Figure 1.**
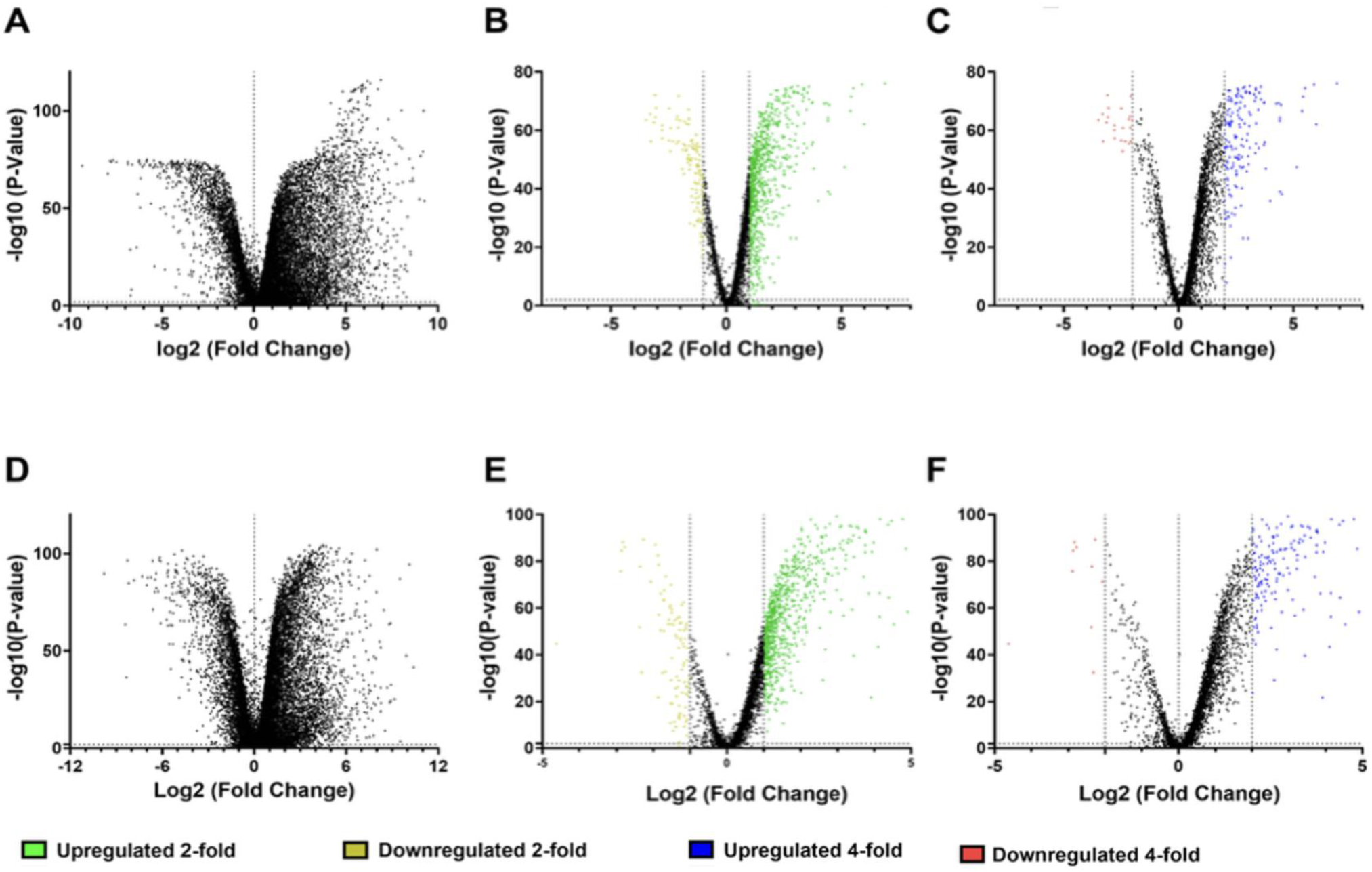
Differential expression landscapes in esophageal (upper, fig. A, B and C) and gastric (lower, fig. D, E and F) tumors. Volcano plots illustrate RNA-seq differential expression (tumor vs. normal) for esophageal (upper panels) and gastric (lower panels) cancers, encompassing 21,479 analyzed genes. The x-axis represents log_2_ fold change (tumor/normal) and the y-axis –log_₁₀_ adjusted p value. (A and D) Global profiles reveal a broader transcriptional activation in esophageal tumors compared with a more balanced pattern in gastric tumors. (B and E) Using a 2-fold cutoff (|log__2__FC| ≥ 1; FDR < 0.05), green dots denote genes upregulated ≥2-fold and yellow dots those downregulated ≥2-fold, showing that esophageal tumors contain a larger fraction of upregulated genes. (C and F) With a 4-fold threshold (|log__2__FC| ≥ 2; FDR < 0.05), blue dots indicate genes upregulated ≥4-fold and red dots those downregulated ≥4-fold, highlighting a subset of strongly induced transcripts in esophageal cancer.

### Identification of genes through condensate biological related functions

To identify biomolecular condensate–associated proteins, we curated a panel of Gene Ontology (GO) terms representing biological processes and subcellular compartments mechanistically linked to condensate formation and function, as described in the material and method section. As displayed in Supplementary Table 1, in esophageal cancer, the most enriched category was positive regulation of transcription by RNA (GO:0045944), encompassing 82 genes including *EPCAM, CREB3L1, MET, SOX4, MYBL2,* and *STAT1*. Other significantly represented terms, such as centrosome, nucleolus, nuclear speckle and PML body, underscored the reorganization of nuclear and RNA-processing domains. Similarly, gastric tumors showed enrichment in positive regulation of transcription by RNA (95 genes; *SOX4, STAT1, MYBL2, FOXM1, MET, VDR, IGF2*), along with nuclear speckle and centrosome, pointing to enhanced transcriptional and mitotic regulation (Supplementary Table 1). In both tumor types, pathways associated with chromatin remodeling, RNA splicing, signal transduction, ubiquitin-dependent proteolysis, stress granule formation, and condensate structures such as P-body and Cajal body were upregulated. Notably, genes such as *TOPBP1* and *CHERP*, both implicated in DNA damage signaling and RNA splicing within phase-separated nuclear condensates assemblies, were significantly upregulated, reinforcing the link between condensate dynamics and genome integrity control (Supplementary Table 1).

Functional enrichment analysis of downregulated genes across esophageal and gastric tumors uncovered a convergent suppression of transcriptional regulatory programs, RNA processing networks, and condensate-associated nuclear compartments **(Supplementary table 2).**

### Condensate-Associated Gene Dependencies Reveal Core Vulnerabilities and Context-Selective Regulators in Esophageal and Gastric Carcinomas

Having identified genes that are recurrently upregulated across tumors, and that can be categorized within protein functions related to biocondensate formation, next we aim to evaluate the functional dependency of the selected candidates. With this approach we aim to select only those with clear role in oncogenesis. Across both esophageal and gastric carcinomas, functional dependency analyses with CRISPR and RNAi screens, as described in the Materials and Methods Section, defined in both tumor types, *HCFC1, TOPBP1,* and *CHERP* as common essential genes (Fig. 2). These factors localize to nuclear condensates such as PML bodies, nuclear specks, and the nucleolus, where they coordinate transcriptional activation, DNA replication stress responses, and RNA splicing. Their depletion strongly impairs tumor cell viability, highlighting them as potential core condensate vulnerabilities in upper gastrointestinal cancers. A second group of genes—including *NCOR1, NCOR2, MYBL2, ATRX, IRS2, ZC3H13, AHR, BMPR2, MAP3K2*, and *WNK2*—were classified as strongly selective (Fig. 2), showing dependency patterns that vary across cell lines, as classified by DepMap^13^. Many of these also participate in condensate-associated mechanisms, such as chromatin remodeling (*ATRX*), transcriptional repression (*NCOR1/2*), or signal-regulated phase transitions (*AHR, MYBL2*). In contrast, genes such as *MED1, MED12, MED13, CDK12, and SCAF11* display no essential dependency, suggestingauxiliary rather than core survival roles (Fig. 2). Together, these findings define a two-tier condensate dependency model in esophageal and gastric cancer: (1) a small, conserved set of nuclear condensate organizers (*HCFC1, TOPBP1, CHERP*) essential for cell viability, and (2) a broader, context-sensitive group that modulates transcriptional plasticity and stress adaptation, supporting tumor evolution rather than baseline survival.

**Figure 2.**
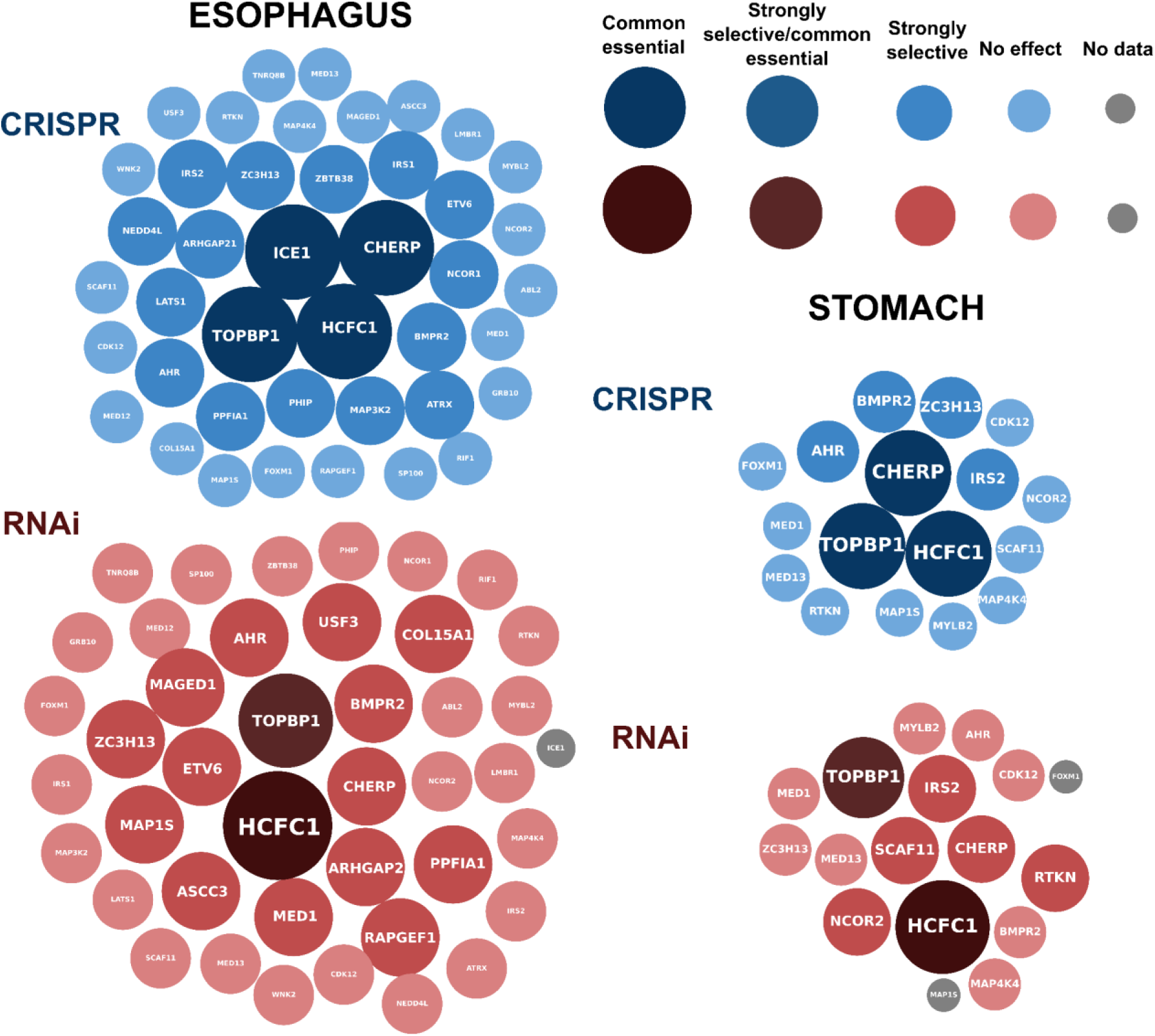
Functional dependency of condensate-associated genes in esophageal and gastric cancers. Effects of CRISPR and RNAi-mediated silencing of genes encoding proteins likely involved in biomolecular condensate formation on the survival of various tumor cell lines. Data depicted were extracted from DepMap portal. Each gene is classified according to its dependency score: common essential (huge), strongly selective/common essential (large), strongly selective (big), no effect (medium), and no data (small). In esophageal tumors, genes such as HCFC1, TOPBP1, and CHERP exhibit strong or common essentiality patterns, indicating their central role in replication stress and transcriptional condensate function. In contrast, gastric cancer cells display a narrower set of dependencies, with TOPBP1, CHERP, and MYBL2 maintaining selective essentiality. Common essential: A gene which, in a large, pan-cancer screen, ranks in the top depleting genes in at least 90% of cell lines, being essential for the survival of most cell lines tested. Strongly selective: A gene whose survival dependency profile is specific to a certain number of tumor cell lines tested. Strongly selective/ common essential: Genes that are essential for survival across most tumor cell types tested but have a special strong effect on survival of a specific subset of tumor cells.

### Expression of condensate-related genes across tumors

To determine whether these candidate condensate-associated dependencies exhibit tumor type–specific or broadly conserved expression programs, we examined their transcriptional profiles across diverse solid malignancies. Therefore, we explored the expression of the previously identified gene across all solid tumors using data from TCGA. Core condensate genes—*HCFC1, TOPBP1,* and *CHERP*—showed uniformly high expression across multiple malignancies, including esophageal, gastric, breast, colorectal, liver, and uterine carcinomas (Fig. 3 A, B). These factors are identified as common essential genes in both esophageal and gastric tumor models, reflecting their indispensable role in sustaining tumor cell viability.

**Figure 3.**
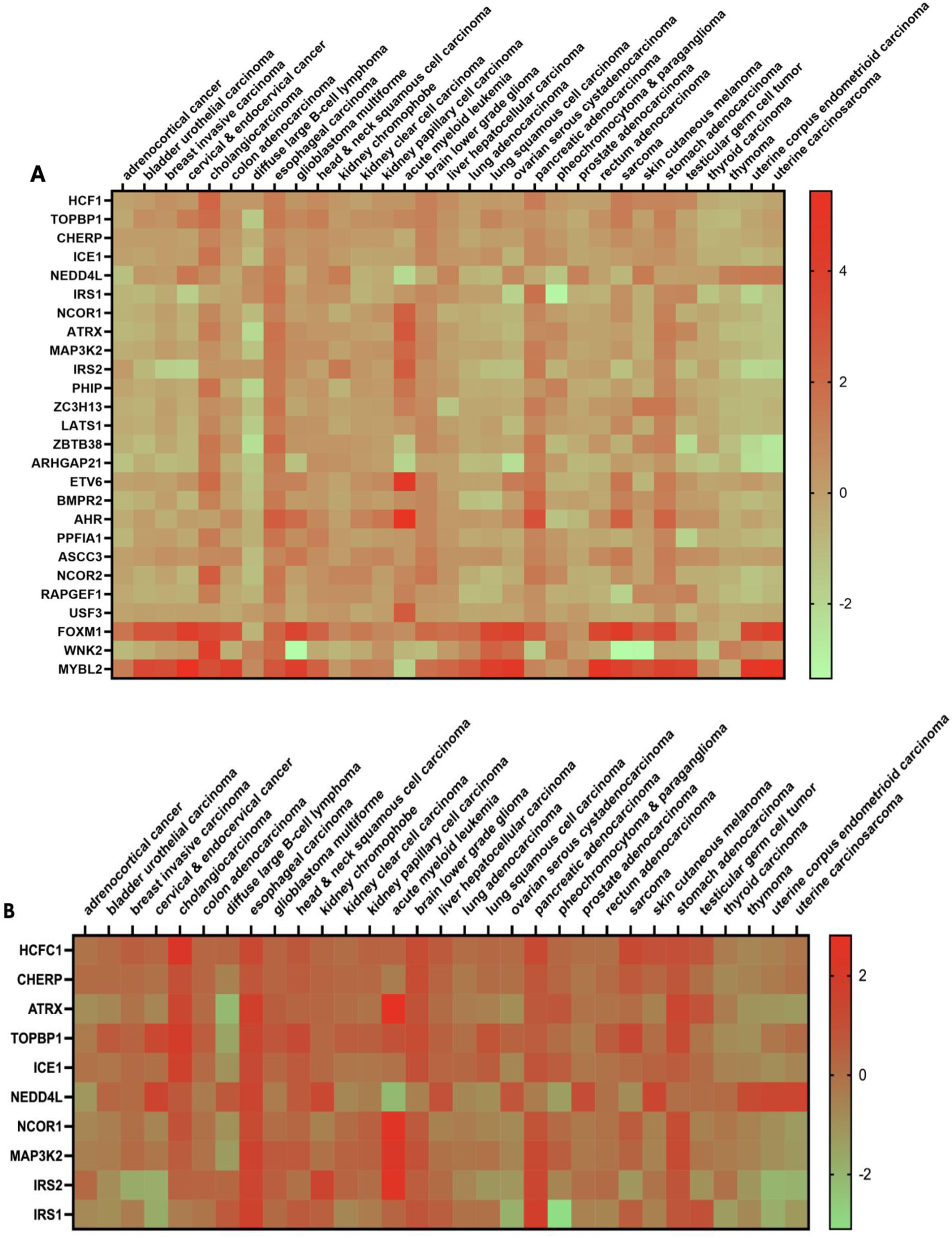
Cross-cancer expression patterns of condensate-associated genes in esophageal and gastric tumors. Heatmaps display normalized expression (log_2_ fold change) of condensate-associated genes across multiple tumor types from TCGA datasets. The **upper panel (A)** corresponds to **esophageal carcinoma**, and the **lower panel (B)** to **gastric cancer**. Each column represents a distinct cancer type, and each row a condensate-related gene. Color intensity reflects relative expression, ranging from downregulation (white) to strong upregulation (dark blue). Esophageal tumors exhibit broader and higher expression levels of condensate regulators—including **TOPBP1**, **CHERP**, **HCFC1**, and **MYBL2**—compared with gastric tumors, which display a more restricted transcriptional activation pattern.

A subset of genes that displayed tumor-selective enrichment including *MYBL2, FOXM1*, and *NCOR2* were prominently expressed in gastric, breast, and uterine carcinomas, where they drive cell-cycle progression and oncogenic transcriptional amplification (Fig. 3 A, B). *NCOR1, ATRX,* and *MAP3K2* were upregulated in esophageal, glioma, and hepatic cancers, implicating them in chromatin remodeling and stress signaling. Meanwhile, *AHR, BMPR2, ZC3H13,* and *IRS2* showed variable expression, reflecting context-dependent condensate engagement (Fig. 3 A, B).

### Biophysical condensate analysis of identified genes

Motivated by the enrichment of condensate-associated genes among upregulated transcripts in both esophageal and gastric tumors, we next applied a more stringent, sequence-based analysis to assess whether the encoded proteins harbor intrinsic features predictive of phase separation. To this end, we estimated the saturation concentration (C_sat_ in μM)—a quantitative measure of phase-separation propensity—using a previously-validated sequence-based ML predictor restricted to IDRs at 293 K and 150 mM of NaCl ^61,67,68,70^. Since many encoded proteins contain a predominant proportion of their residues within globular domains, we restricted our analysis to canonical sequences with at least 40% of residues within IDR segments across the entire sequence, as determined using AlphaFold3 confidence scores (pLDDT < 70; see Materials and Methods) ^63^. The filtered IDR-rich proteins were subsequently evaluated using the ML predictor to obtain their corresponding C_sat_ values (**Fig. 4**). For multidomain proteins containing multiple IDRs interspersed with different globular domains (GD), e.g. IDR1–GD1–IDR2–GD2–IDR3, individual disordered segments were concatenated (e.g., IDR1–IDR2–IDR3) and supplied as a single sequence to the ML predictor, thereby estimating the aggregate IDR-driven C_sat_ while excluding folded domains Archetypal phase-separating proteins, such as FUS ^71,72^, hnRNPA1^39^, or TDP-43^73,74^) typically exhibit experimental saturation concentrations in the low micromolar range (∼1 to 10μM ^38,41,72,75^) and contain well-defined IDRs. Accordingly, we considered proteins with predicted C_sat_ values below ∼20 μM and substantial IDR content as strong candidates for intrinsic condensate formation (dark red shadowed area in **Fig. 4**, please see Supp Table 3 for the predicted value for all the sequences). Consistent with experimental data, the control phase-separating protein FUS (red star) localized within this region, with a total of 480 amino acids in IDRs and a predicted C_sat_ of ∼1μM, closely matching experimental values(1–5 μM ^72,76^). More importantly, in both types of tumors, proteins such as *TOPBP1, CHERP,* and *AHR* among few others cluster within the low C_sat_ region (i.e., below 10 μM) and display extensive IDRs content (300–700 residues), consistent with phase-separation propensities comparable to established condensate-forming proteins. Crucially, only *TOPBP1* has been previously identified as an essential regulator in transcriptional condensates and replication stress responses, supporting their potential roles as structural and functional components of biomolecular condensates. A second tier of candidates includes proteins with extremely high numbers of residues within disordered regions (above 1000 residues in IDRs), such as *HCFC1, IRS2, NCOR1,* or *NCOR2* among others, which also exhibit relatively low predicted C_sat_ values. Nevertheless, beyond their sequence length and molecular weight, phase-separation propensity is strongly influenced by residue composition and sequence patterning^77^. Finally, proteins such as *MYBL2* and *FOXM1* lie near the predicted phase-separation threshold, with C_sat_ values between 10–100 μM (comparable to the experimental values for FUS-LCD) ^76^, suggesting that they require cooperative interactions or additional partners to form condensates.

**Figure 4.**
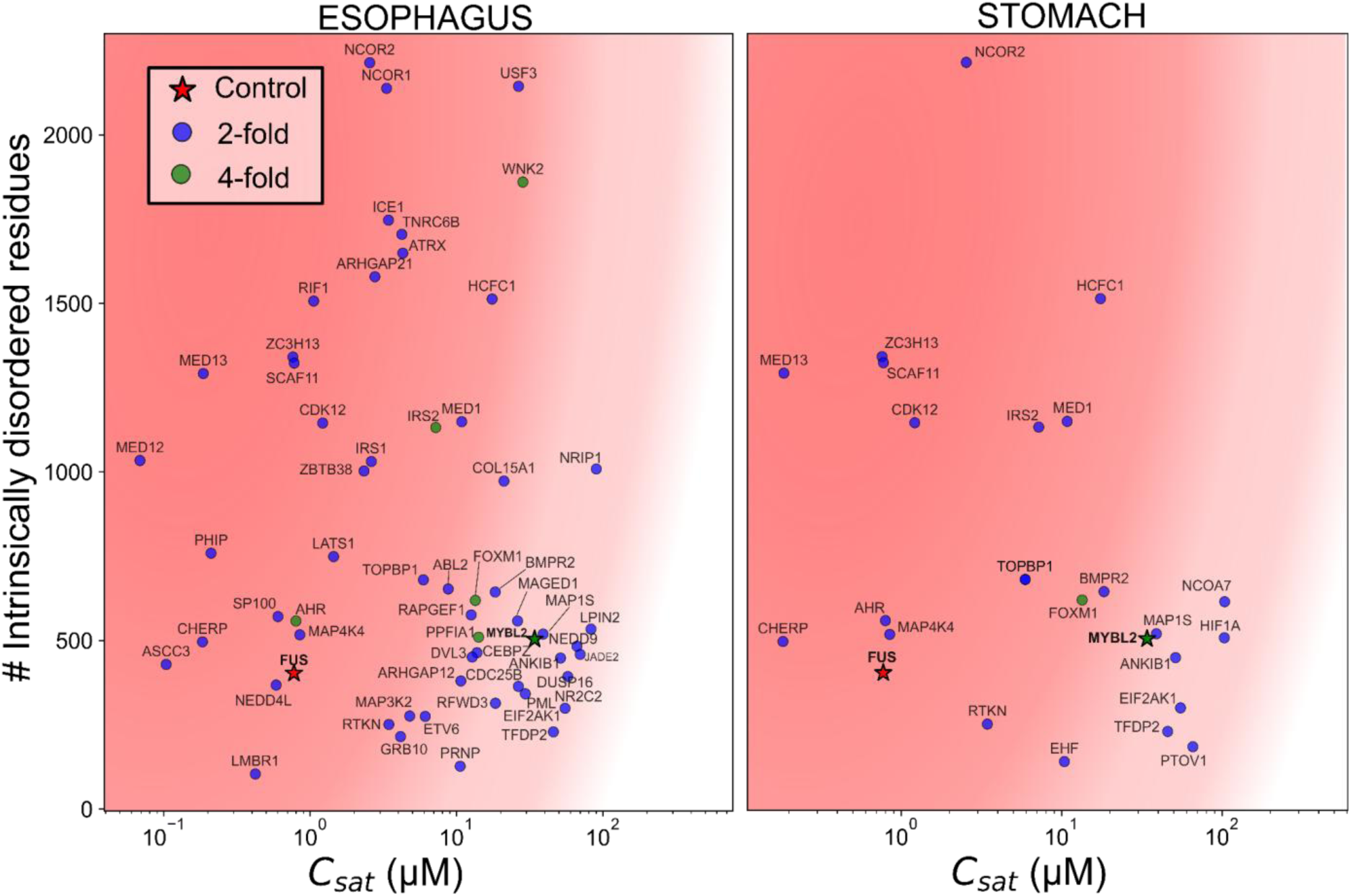
Phase separation prediction in esophageal (left) and gastric (right) tumors. Map plot of different proteins involved in esophageal (left panel) and gastric (right panel) cancers representing the number of residues in intrinsically disordered regions (IDRs) as a function of the predicted saturation concentration C_sat_ in *μ*M using the CALVADOS2 ML predictor^61^. We include data for 2-fold (blue) and 4-fold (green)e proteins as well as a control protein (FUS) highlighted in red as an archetypal phase-separating sequence. The red shadowed area represents the regime where condensation is expected (e.g., below 20 *μ*M).

### Condensate formation of selected genes predicted by sequence-dependent molecular dynamics simulations

To determine whether the predicted IDR-encoded phase-separation propensities give rise to emergent condensate behavior in full-length multidomain proteins involved in esophagus and gastric cancer, we performed MD simulations with the Mpipi-Recharged model. Mpipi-Recharged is a state-of-the-art sequence-dependent coarse-grained model which has been shown to quantitatively reproduce phase behavior and material properties across diverse biomolecular condensate systems ^64,66–68,73^ (see Materials and Methods section for further details on the model potential). We carried out DC simulations^78^ of solutions of CHERP, TOPBP1, HCFC1, and IRS2, which were selected based on their predicted phase-separation propensity (Fig. 5) and prior classification as essential or selectively expressed genes. (Fig. 4). Folded domains were now defined as regions with pLDDT > 90 from AlphaFold predictions and modeled as rigid bodies (**Fig. 5A**, see Methods for details). Our simulations are performed at 280K and 150 mM NaCl.,As previously reported^66,73^, Mpipi-Recharged exhibits a maximum offset of ∼20–30 K in predicted critical solution temperatures^64,66^. Therefore, proteins which exhibit phase-separation at 280K are expected to form condensates at physiological conditions (e.g., ∼310K)). Both CHERP and TOPBP1 displayed clear phase separation behaviour in DC simulations (stable slab formation, sharp interfaces, and distinct coexistence densities between dilute and condensed phases; **Fig. 5B**), *CHERP,* in particular, formed a stable condensed phase with an approximate density of ∼0.4g/cm^3^, comparable to values reported for FUS and hnRNPA1-LCD ^66^ under the same conditions. These results are consistent with their low predicted Csat values (**Fig. 4**). In contrast, IRS2 and HCFC1 remained homogeneously distributed across the simulation box forming a homogeneous solution, showing no evidence of spontaneous phase separation (**Fig. 5B**). Although their predicted C_sat_ value (C_sat_∼10μM, **Fig. 4)** suggested potential phase-separation propensity, our Mpipi-Recharged simulations indicate that their specific residue composition and sequence patterning, together with competition between intermolecular IDR–IDR contacts and intramolecular IDR–globular interactions, limit the formation of a sufficiently connected multivalent network under the conditions analyzed. These results further indicate that C_sat_ values derived from IDR-only predictors must be carefully interpreted in the context of full-length proteins, as folded domains and intramolecular interactions can substantially modulate effective phase-separation propensity.

**Figure 5.**
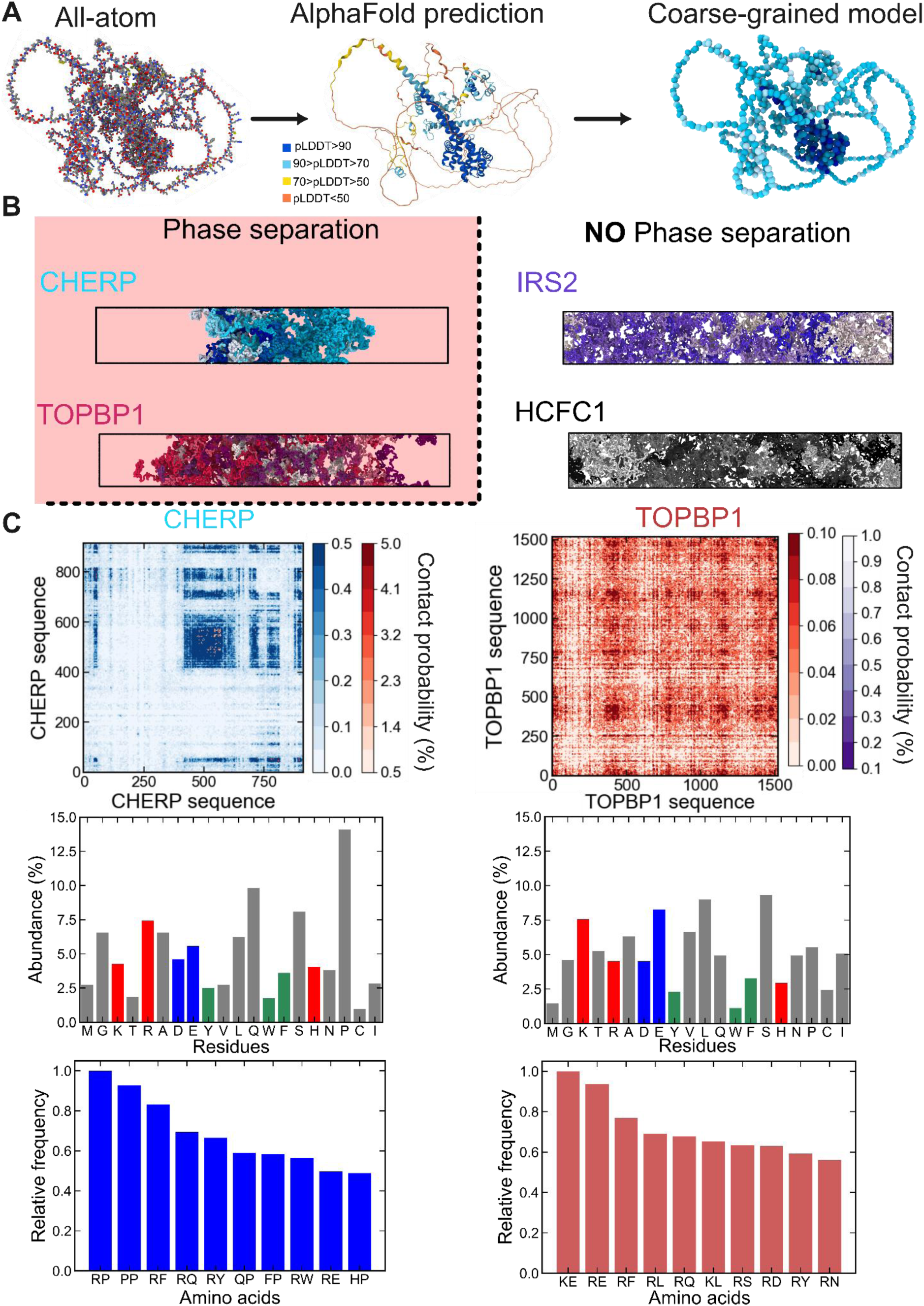
Molecular simulations of TOPBP1 and CHERP reveal their condensation propensity. **A.** Modelling approach to build the coarse-grained representation of CHERP. We use the position of the all-atom model and the prediction of Alphafold to distinguish between globular (pLDDt>90) and intrinsically disordered domains (pLDDt<90). The CG model represents the IDRs in skyblue and the globular domains in dark blue, where each bead represents one amino acid. **B.** Representative snapshots from direct coexistence simulations of CHERP, TOPBP1, IRS2, and HCFC1 at 280 K and 150 mM of NaCl. The simulations of CHERP and TOPBP1 clearly demonstrate two distinct phases while HCFC1 and IRS2 do not phase-separate. We have depicted each protein replica in a different color for all the presented systems. **C. Top**: Intermolecular contact frequency for CHERP (left) and TOPBP1 (right) from our DC simulations. The colorbar shows the probability of residue-residue contacts along the sequence of both proteins, from 0-0.5% and 0.5-5% for CHERP, and from 0-0.1% and 0.1-1% for TOPBP1, given the probability of contact for each protein replica in the system. **Centre**: Amino acid composition of CHERP (left) and TOPBP1 (right) in abundance (%). We have coloured positively charged residues in red, negatively charged residues in blue, and aromatic amino acids in green due to their relevance in biomolecular condensation. **Bottom:** Most frequent pairwise residue-residue interactions from our simulations and normalized by the maximum number of contacts.

To further elucidate the mechanisms and molecular interactions that drive *CHERP* and *TOPBP1*condensate formation, we analyzed intermolecular residue–residue contact maps from DC simulations (**Fig 5C**). Contacts were defined as the time-averaged fraction of residue pairs within the condensate closer than a cutoff distance, corresponding to attractive interactions in the coarse-grained model (see Methods for details) ^79^. These contacts quantify the intermolecular connectivity that stabilizes the condensed phase and enable identification of the sequence regions contributing most strongly to multivalent network formation.

*CHERP* comprises three different IDR regions (IDR1: residues 1–13; IDR2: 21–123; IDR3: 327–916) separated by two globular domains (GD1: 14–20; GD2: 124–326). Its intermolecular contact map reveals an interaction hotspot within IDR3, spanning residues 450–600, which constitutes the dominant region driving intermolecular connectivity of the *CHERP* condensates Crucially, the identification of this interaction hotspot indicates that CHERP condensate stabilization is most strongly driven by a limited sequence segment, establishing a rational entry point for therapeutic disruption of the ensuing condensates. Targeted modulation of this segment could selectively inhibit condensate formation and thereby influence condensate-dependent tumorigenic processes.

Detailed analysis of the most frequent pairwise contacts in *CHERP* condensates (**Fig 5C, left bottom**) reveals that arginine (R) plays a central role in condensate stabilization, primarily establishing contacts with proline (R–P) and cation–π interactions with aromatic residues (R–F and R–Y). Importantly, the high abundance of proline in *CHERP* (near 15% in **Fig 5C, left centre**), compared to aromatic residues (<5%), clarifies the prevalence of R–P contacts. These findings are consistent with the dominant contribution of arginine-mediated interactions, including cation–π contacts with aromatic residues, to the multivalent network stabilizing condensates involving RNA-binding proteins^80–82^.

When the same analysis is applied to *TOPBP1* (**Fig. 5C, right**), we found a more homogeneous contact map featuring a slightly more pronounced binding domain between residues 400 and 500. Importantly, the maximum contact probability per residue in *TOPBP1* is considerably lower than that observed for *CHERP* (1% compared to 5%), evidencing its lower propensity for phase separation, and the lower condensate protein density at the studied conditions (0.20 vs 0.4 g/cm^3^). Analysis of the most frequent amino acid pairwise interactions (**Fig. 5C, bottom right**) underscores the importance of electrostatic contacts, primarily between lysine or arginine and glutamic acid (KE and RE pairs), in mediating *TOPBP1* condensation. Consistently, **Fig. 5C** reveals that charged residues (R, K, D, and E) constitute approximately 25% of the total *TOPBP1* sequence. Electrostatic interactions play a critical role in regulating condensate formation, with minor spontaneous variations in intracellular salt gradients capable of markedly disturbing the balance between condensate stability and dissolution ^83,84^. Moreover, cation-𝜋 interactions, such as those established between arginine with both tyrosine and phenylalanine, also constitute a major LLPS-stabilizing binding mode for TOPBP1.

Since it has been already reported that TOPBP1 is capable of forming condensates under physiological conditions^85^—fully consistent with our computational predictions (Fig. 6)—but there is no experimental evidence of CHERP undergoing phase-separation, we performed turbidity assays of CHERP at room temperature to estimate its saturation concentration and directly compare with our ML predictions and Mpipi-Recharged simulations (**Fig. 6A)**. The experimental value for the saturation concentration of CHERP is ∼0.07mg/mL, equivalent to 0.7μM. Such saturation concentration (C_sat_ ≈ 0.7 μM) falls within the typical range reported for canonical scaffold proteins that readily form condensates (e.g., FUS: ∼2-4 μM ^72^, TDP-43: ∼8 μM^74^, hnRNPA1: ∼15μM^39^). This value is also consistent with the predicted C_sat_ by the CALVADOS3 ML algorithm (C_sat_ ≈ 0.2 μM, **Fig. 5** and inset in **Fig. 6A**). Moreover, the experimental C_sat_ is in good agreement with our DC simulations estimate using the Mpipi-Recharged (C_sat_ < 2 μM) which yielded a phase diagram showing phase-separation over a wide range of temperatures (**Fig. 6B**). We acknowledge that obtaining a fully rigorous estimate of C_sat_ directly from our DC simulations is computationally demanding for a sequence of this length, as an accurate determination would require substantially longer DC simulations—i.e., at least one order of magnitude longer simulations to ensure significant sampling of the dilute phase, or alternatively the application of a sophisticated thermodynamic integration framework^66^. Nevertheless, we have estimated an approximate upper bound of C_sat_ within our simulation timescale (∼2 μs) of approximately C_sat_ <1.9 μM at 310K from the coexistence concentrations obtained in our DC simulations (black dashed line in **Fig. 6A**). This magnitude has been estimated by computing the average number of events in which a given protein replica within the simulation trajectory (i.e. protein probability) is present at the volume of the dilute phase. Furthermore, we found that the critical solution temperature of CHERP (**Fig. 6B**; T_c_∼365K) is similar to those of several aromatic variants of hnRNPA1-LCD^38,66,68^ which also displayed similar values of T_c_ (e.g., within the range of 320 to 360K) and C_sat_ values from 1 to 10 μM^38,66,86^. Moreover, other archetypal phase-separating proteins such as FUS (T_c_∼330K)^54^ or TDP-43 (T_c_∼305K)^73^ also displayed C_sat_ values between 1 and 15 μM at physiological conditions, both experimentally, and in silico using the Mpipi-Recharged model^66–68^. In **Fig. 6B**, we also included the full phase diagram of CHERP in the temperature-density plane as well as simulation snapshots at two different temperatures (as indicated by the horizontal lines), with the region spanning amino acids 450–600—shown in magenta—highlighted for its high contact frequency (**Fig. 5C**) and its primary role in sustaining phase-separation.

**Figure 6.**
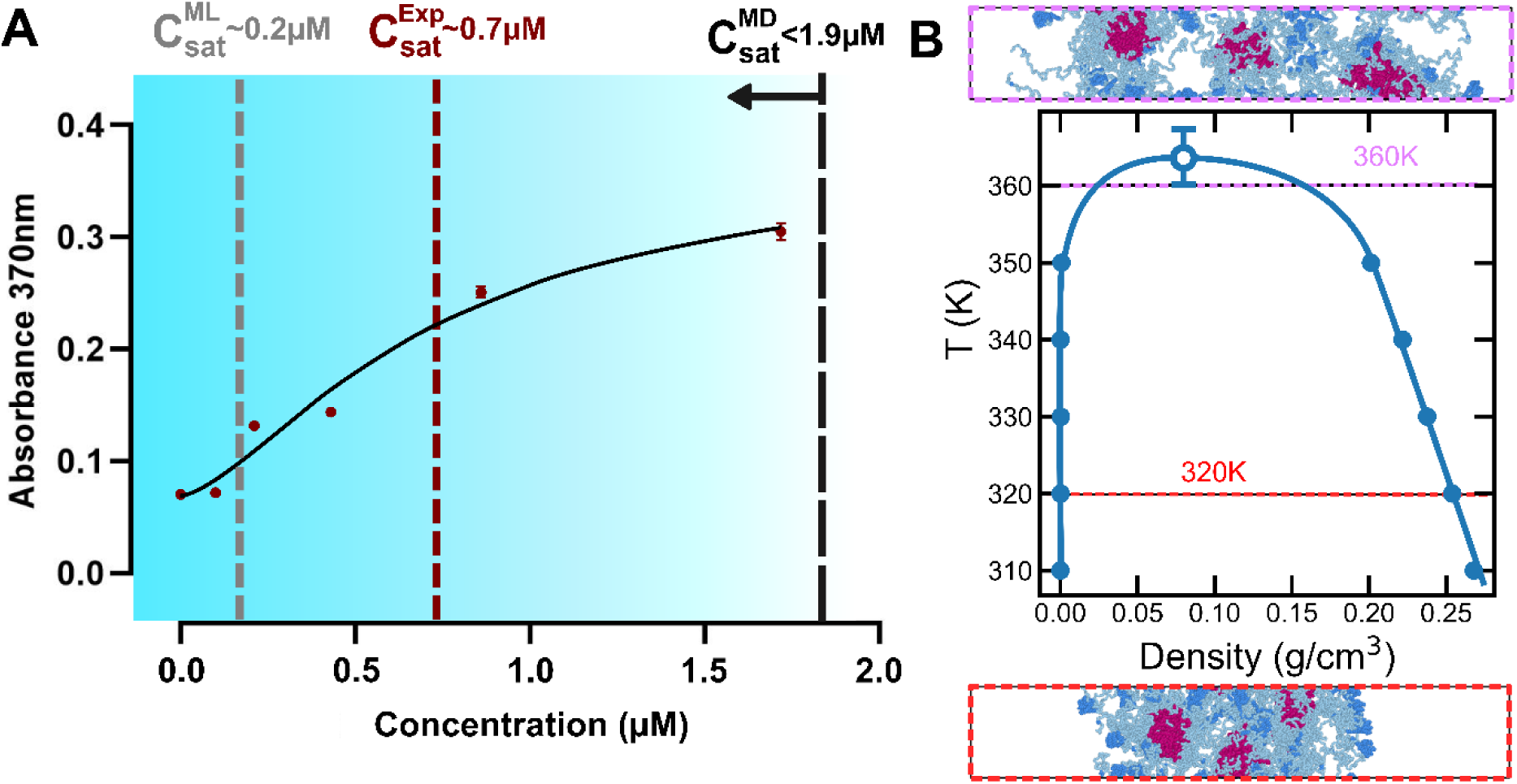
Experimental turbidity assays and molecular simulations predict condensate formation of CHERP. **A.** Determination of CHERP saturation concentration from a turbidity assay (room temperature, 150 mM NaCl). A sigmoidal fit to the data was used to calculate the value, indicated by the dotted vertical lines. Dashed lines indicate the position of the C_sat_ value from the ML predictor (gray) and from our experiments (dark red). Light blue gradient and black dashed line indicate the area of the expected C_sat_ value (C_sat_<1.9μM) from our DC simulations. **B.** Phase diagram of CHERP in the temperature-density plane from DC simulations. The critical point was calculated with the law of rectilinear diameters and critical exponents. Top and bottom snapshots represent typical configurations from our simulations colouring the globular domains in dark blue and highlighting the 450-600 amino acids in CHERP sequence in magenta.

## Discussion

This study reveals that esophageal and gastric cancers share a condensate-enriched transcriptional architecture characterized by the activation of proteins containing large IDRs which drive phase-separation, and coordinate transcription, RNA processing, and stress adaptation. By integrating transcriptomic, functional, and biophysical data, we demonstrate that a subset of nuclear regulators—including *TOPBP1* and *CHERP*—constitute a conserved condensate core that sustains tumor cell viability. Our findings expand the emerging concept that cancer progression involves not only genetic and epigenetic alterations but also a fundamental reorganization of the cell’s mesoscale architecture through biomolecular phase-separation^13^

Our analyses identify *TOPBP1* and *CHERP* as key components of condensate-like molecular architectures. *TOPBP1* (Topoisomerase II Binding Protein 1) is a scaffold for DNA replication and damage-response condensates, where it interacts with ATR and 9-1-1 complexes to coordinate replication stress checkpoints ^87^. Its intrinsically disordered N-terminal region and multivalent BRCT domains enable condensation at stalled replication forks and DNA lesions ^85^. Similarly, *CHERP* (Calcium Homeostasis Endoplasmic Reticulum Protein) localizes to nuclear specks and spliceosomal condensates, where it contributes to pre-mRNA splicing and calcium-regulated transcriptional programs ^88^. Both proteins exhibit large intrinsic disorder regions and low predicted saturation concentrations, supporting strong biophysical propensity for phase separation. Functionally, they emerge as essential genes in CRISPR and RNAi dependency datasets, suggesting that upper gastrointestinal cancers rely on their condensate scaffolding functions to maintain transcriptional and replication fidelity under oncogenic stress.

These results align with the growing recognition that condensate biology provides a structural and kinetic framework for oncogenic transcriptional amplification ^89^. Oncogenic transcription factors, coactivators, and chromatin remodelers—such as *MYC, MED1*, and *BRD4*—assemble into transcriptional condensates that concentrate polymerases and epigenetic modifiers at super-enhancers, thereby sustaining high transcriptional throughput ^90^. Our study extends this paradigm to esophageal and gastric cancers, demonstrating that these tumors harbor a condensate-driven transcriptional state reinforced by upregulation of genes governing chromatin remodeling, RNA splicing, ubiquitin-dependent proteolysis, and signal transduction. The enrichment of nuclear speckles, PML bodies, nucleoli, and centrosome components indicates a global remodeling of phase-separated compartments, consistent with previous reports linking condensate reorganization to proliferation and therapy resistance ^13^.

We identify two tiers of condensate dependency. The first consists of conserved nuclear scaffolds—HCFC1, TOPBP1, and CHERP—required for tumor cell viability and organization of transcriptional and replication-stress condensates^13^. The second includes context-dependent regulators (MYBL2, FOXM1, NCOR1/2, ATRX, AHR, MAP3K2) with lineage-specific expression and selective dependencies ^91^.

Very recently, *TOPBP1* condensation has been recognized as an essential amplifier of ATR signaling that enables cancer cells to endure replication stress. Pharmacologic disruption of this process by AZD2858 abrogates TOPBP1 self-interaction and ATR activation, intensifying DNA damage and apoptosis ^89^. In colorectal cancer models, AZD2858 synergizes with FOLFIRI chemotherapy to suppress tumor growth^78^. These findings unveil condensate disassembly as a tractable vulnerability within the replication-stress response network^90^. To the best of our knowledge, no study has reported yet the role of *CHERP* in condensate biology.

The integration of gene-expression magnitude, phase-separation propensity, and functional dependency represents a distinctive aspect of our approach. Traditional genomic analyses often identify driver alterations without capturing how these alterations reconfigure subcellular organization^91^. Here, combining transcriptomic data with phase-separation predictors and high-resolution sequence-dependent coarse-grained simulations provides an additional biophysical layer that contextualizes oncogenic dependencies. For instance, *TOPBP1* and *CHERP* cluster in a biophysical regime of large IDRs which enable low C_sat_ values to undergo phase-separation, whereas *HCFC1, IRS2, NCOR1/2* and *ATRX* occupy an intermediate regime indicative of conditional condensation upon partnering with additional scaffolds. More importantly, our turbidity assay has confirmed the extraordinary propensity of CHERP to form condensates, in agreement with our Mpipi-Recharged simulations and the CALVADOS3 ML predictor. This mapping suggests that cancer cells exploit a continuum of condensate behaviors, from constitutive structural scaffolds to tunable, stress-responsive assemblies. The functional implications of condensate reprogramming are diverse as condensates concentrate macromolecules, accelerate enzymatic reactions and buffer protein fluctuations^92^.

From a translational standpoint, our findings suggest that targeting condensate integrity may open a new therapeutic dimension in upper GI cancers. Several strategies can be envisioned. Small molecules or peptides that alter protein valency, disrupt IDR-mediated interactions, and/or modulate local charge distribution could selectively dissolve oncogenic condensates^23^. Similarly, inhibitors of post-translational modifications that regulate condensate stability—such as phosphorylation or acetylation—could indirectly modulate condensate dynamics ^23^. In this context, *TOPBP1* and *CHERP* represent particularly attractive targets, as their essential condensate scaffolding roles are conserved yet largely tumor-restricted in their overexpression.

Biophysical screening platforms integrating phase-separation assays with ML-based predictors and MD simulations, such as the one described here, may accelerate the discovery of condensate-modulating compounds with therapeutic potential. In summary, using an integrative computational framework, we uncover a shared condensate-based transcriptional network anchored by *TOPBP1* and *CHERP*, whose essentiality defines a core condensate dependency in these malignancies. This condensate-driven model not only advances our understanding of tumor cell organization but also delineates a new frontier for therapeutic innovation in upper gastrointestinal oncology.

## Acknowledgements

A. R. T. acknowledges funding from the European Union Horizon 2020 research and innovation programme (grant agreement 803326 to R.C.-G.) and from Ministerio de Ciencia e Innovacion under the Juan de la Cierva fellowship (JDC2024-053759-I). R.C.-G. acknowledges funding from the European Research Council (ERC) under the European Union Horizon 2020 research and innovation programme (grant agreement 803326). J. R. E. acknowledges funding from Emmanuel College, the University of Cambridge, the Ramon y Cajal fellowship (RYC2021-030937-I), the Spanish scientific plan and committee for research reference PID2022-136919NA-C33, and the European Research Council (ERC) under the European Union’s Horizon Europe research and innovation program (grant agreement no. 101160499). A.O. and C.P. acknowledge funding from CRIS Cancer Foundation (AOF.C01CRIS and AOF.M01CRIS). This work has been performed using resources provided by the Cambridge Tier-2 system operated by the University of Cambridge Research Computing Service (http://www.hpc.cam.ac.uk) funded by EPSRC Tier-2 capital grant EP/P020259/1-CS170. This work has also been performed using resources provided by Archer2 (https://www.archer2.ac.uk/) funded by EPSRC Tier-2 capital grant EP/P020259/e829. The authors also thankfully acknowledge RES computational resources provided by Mare Nostrum 5 through the activity 2024-3-0001.

