## Supplementary Material for "Condensate-Driven Transcriptional Reprogramming Defines Core Vulnerabilities in Esophageal and Gastric Cancers"

**A**

| GO Term | GO ID | Category | Number of genes | Representative genes | p-value adjusted |
|---|---|---|---|---|---|
| Positive regulation of transcription by RNA | GO:0045944 | BP | 82 | EPCAM,CREB3L1,MET,SOX4,LUM,MYBL2,ITGA6,AGRN,MAVS,SOX9,NR2C2,SLC40A1,TOP2A,STAT 1,FOXM1,HELZ2,AHR,F2RL1,HLTF,ZFHX3,LRP5 | $9.330\times10^{-96}$ |
| Centrosome | GO:0005813 | CC | 43 | SERINC5,ECT2,CDK6,CENPF,CCDC88A | $4.696\times10^{-59}$ |
| Nucleolus | GO:0005730 | CC | 56 | RPS17,GPRC5A,KRT18,TOP2A,STAT1,ITGB4,CEMIP2,HLTF | $1.160\times10^{-46}$ |
| Signal transduction | GO:0007165 | BP | 79 | MET,MDK,PLAU,GPRC5A,PLAUR, AGRN,MAVS,CDK6,SOX9,ALCAM,VAV2,STAT1,PTPRK,PPFIA1,IRS2 | $2.506\times10^{-38}$ |
| Nuclear speck | GO:0016607 | CC | 23 | SRSF10,FTO,LIMK1,ZC3H13,PAK2,TCF12,NRIP1,MED1,CDK12,PRKAA1,CNOT7,PSME4,PIAS1,SP1 00,RNPS1,HIF1A,DHX36,SF3A2,PATL1,ASCC3,NONO,VIRMA,CDC5L | $6.851\times10^{-30}$ |
| Regulation of DNA-templated transcription | GO:0006355 | BP | 39 | ZNF264,SRSF10,NR2C2,SOX4,ZFHX3,AHR | $2.056\times10^{-28}$ |
| PML body | GO:0016605 | CC | 12 | SKIL,ATRX,ELF4,TOPBP1,PML,PIAS1,SP100,SQSTM1,RB1,PATL1,RFWD3,NBN | $1.070\times10^{-26}$ |
| Chromatin remodeling | GO:0006338 | BP | 17 | SOX9,ATRX,HCFC1,ACTL6A,CHRAC1,YEATS2,PML,RSF1,RBBP4, DEK,BAZ1B,MTA1,RB1,PAK1,TOP1,BAZ1A,CBX3 | $1.687\times10^{-22}$ |
| P-body | GO:0000932 | CC | 11 | AGO2,IGF2BP2,PNRC2,TNRC6B,DDX6,CNOT7,SQSTM1,PATL1,CAPRIN1,TNRC6B,TOP1 | $2.720\times10^{-22}$ |
| Ubiquitin-dependent protein catabolic proces | GO:0006511 | BP | 13 | NEDD4L,DTX3L,UBE2C,ANKIB1,MDM2,RNF213,UBE2L6,USP38,MIB1,UBE3A, SQSTM1,PSMD3,PSMD11 | $1.802\times10^{-17}$ |
| Cytoplasmic stress granules | G0: 0010494 | CC | 7 | IGF2BP2,ZNFX1,DDX6,STAU1,PRRC2C,PABPC4,CAPRIN1, | $2.883\times10^{-15}$ |
| Chromatin organization | GO:0006325 | BP | 18 | LOXL2,TOP2A,HLTF,NUCKS1,ATRX,TBL1XR1,ANP32E,RSF1 NCOR1,PRKAA1,TCF3,H2AFV,RBBP4,DEK,ARID4B,RB1,BUD23,PCNA, | $4.339\times10^{-14}$ |
| Transcription by RNA polymerase II | GO:0006366 | BP | 16 | NFE2L3,SOX9,SLC40A1,CEBPB,ATRX,FOSB,IRF1,ETS1,TAF2,POLR2K,CDK12, DEK,PARP1,SQSTM1,ARID4B,RB1 | $7.796\times10^{-12}$ |
| RNA splicing | GO:0008380 | BP | 12 | ZC3H13,MBNL2,USB1,MBNL3,CDK12,SF3B4,SCAF11, RNPS1,SF3A2,NONO,SNRPD1,VIRMA | $2.454\times10^{-10}$ |
| Proteolysis involved in cellular protein catabolic | GO:0051603 | BP | 21 | PSMB9,CTSS,PSMB2,MDM2,CTSZ,CTSC,CTSH,CTSB,CTSL,PSMB8,PKN1,SOCS3,MAP4K4,ROCK1, SQSTM1,RPS6KA3,PRKCSH,DVL3,PAK1,ARFGEF2,TGFBR1 | $6.849\times10^{-8}$ |
| Intracellular signal transduction | GO:0035556 | BP | 18 | MDK,WNK2,ECT2,MAVS,MFHAS,PRKCA,MYADM,LRRC58,TFRC,ERBB2,PAK2,PRKCI,PRNP,LRRC59 ,MYO5A,STK4,ZBTB33,MAP3K2 | $1.405\times10^{-7}$ |
| Cajal body | GO:0015030 | CC | 2 | XPO1,ICE1 | $4.890\times10^{-3}$ |
| RNA catabolic process | GO:0006401 | BP | 3 | OAS2,XRN2,PABPC4 | $1.666\times10^{-3}$ |
| Cytoplasmic ribonucleoprotein granule | GO:0036464 | CC | 2 | PABPC4,IGF2BP2 | $3.086\times10^{-2}$ |
| RNA processing | GO:0006396 | BP | 6 | RBMS2,ADAR, CHERP, XRN2,PABPC4, SNRPD1 | $1.384\times10^{-2}$ |

**B**

| GO Term | GO ID | Category | Number of genes | Representative genes | p-value adjusted |
|---|---|---|---|---|---|
| Positive regulation of transcription by RNA | GO:0045944 | BP | 95 | S100A10, VDR, AGRN, SOX4, STAT1, LUM, KLF5, MAVS, ONECUT2, IGF2, TOP2A, MYBL2, FOXM1, MACC1, AGO2, MET, SOX9, EPCAM, | $2.89\times10^{-106}$ |
| Nuclear speck | GO:0016607 | CC | 47 | SRSF10, SMC4, SRPK1, NEK6, CDK12, MED1, ASCC3, SNRPB2, CARMIL1, LIMK1, PIAS1, ZNF217 | $1.79\times10^{-32}$ |
| Centrosome | GO:0005813 | CC | 23 | SERINC5, CENPF, ECT2, PLK1 | $9.86\times10^{-27}$ |
| Regulation of DNA-templated transcription | GO:0006355 | BP | 35 | HMGA1, VDR, SRSF10, SOX4, LBH, IGF2, CDCA7 | $3.54\times10^{-25}$ |
| P-body | GO:0000932 | CC | 11 | PNRC2, AGO2, DDX6, CAPRIN1, PATL1, TOP1, CNOT7, UBAP2, SAMD4B, PUM1, IGF2BP2 | $1.08\times10^{-24}$ |
| Chromatin remodeling | GO:0006338 | BP | 15 | SOX9, BAZ1A, CHRAC1, HDGF, CHD7, ZBTB7A, ACTL6A, TOP1, DEK, CBX3, REST, HCFC1, DDX21, RUVBL2, RB1 | $1.35\times10^{-19}$ |
| Intracellular signal transduction | GO:0035556 | BP | 26 | ERBB2, MAVS, PPP1R1B, ECT2, MDK | $1.32\times10^{-16}$ |
| RNA splicing | GO:0008380 | BP | 17 | SNRPF, PTBP3, SRPK1, SNRPB, CDK12, MBNL3, SF3B4, ZC3H13, SCAF11, C1QBP, PTBP1, RNPS1, LGALS3, KHSRP, SF3A2, ESRP1, | $1.38\times10^{-15}$ |
| PML body | GO:0016605 | CC | 7 | TOPBP1, RB1, TP53, SKIL, PIAS1, NBN, PATL1 | $8.48\times10^{-15}$ |
| Ubiquitin-dependent protein catabolic proces | GO:0006511 | BP | 10 | UBE2C, DTX3L, MDM2, NDFIP2, RNF213, UBE2L6, PSMD3, ANKIB1, PSMD11, PSMD14 | $6.13\times10^{-13}$ |
| Proteolysis involved in cellular protein catabolic process | GO:0051603 | BP | 10 | PSMB9, CTSZ, CTSH, IDE, CTSC, CTSS, CTSB, PSMB2, MDM2, PSMB8 | $5.86\times10^{-12}$ |
| Transcription by RNA polymerase II | GO:0006366 | BP | 16 | HNF4A, NFE2L3, SOX9, RB1, SLC40A1, CDK12, TFDP1, CBFB, POLR2K, CHD7, DEK, IRF1, DDX21, TRIM24, RREB1, PARP1 | $7.80\times10^{-12}$ |
| Signal transduction | GO:0007165 | BP | 26 | MDK, MET, IL2RG, ERBB2, SOX9, CEACAM1, FAM83H, GPRC5A, AGRN, STAT1, MAVS, PLAU, OLFM4, PPP1R1B | $7.83\times10^{-05}$ |
| Cajal body | GO:0015030 | CC | 2 | DKC1, NOP58 | $4.89\times10^{-03}$ |
| Cytoplasmic stress granules | G0: 0010494 | CC | 8 | IGF2BP2, PABPC1, DDX6, ZNFX1, CAPRIN1, PRRC2C, KHSRP, PUM1 | |
| Chromatin organization | GO:0006325 | BP | 1 | TOP2A, CHD7, ZBTB7A, TCF3, DEK, HMGB2, TBL1XR1, BANF1, PCNA, LOXL2, RB1, ANP32E, NUCKS1 | |
| RNA processing | GO:0006396 | BP | 5 | SNRPD1, RBMS2, DKC1, CHERP, ADAR | |

*Supplementary Table 1.* ***Gene Ontology (GO) enrichment analysis of upregulated genes in esophageal (A) and gastric (B) tumors.*** *List of significantly enriched GO terms (FDR < 0.05) among genes upregulated in esophageal cancer compared with normal tissue. The table shows the GO term name, GO identifier (GO ID), functional category (BP = biological process; CC = cellular component), number of genes contributing to each term, representative genes within each category, and the corresponding adjusted p value.*

**A**

| GO Term | GO ID | Category | Number of genes | Representative genes | p-value adjusted |
|---|---|---|---|---|---|
| Nucleolus | GO:0005730 | CC | 13 | SRSF5,PIM1,S100A16,CSTB,PLEKHM1,RPS6,STK24,NOP53,DNAJB1,SFN,RPS10,RPL26,ACADVL | $3.561\times10^{-10}$ |
| Nuclear speck | GO:0016607 | CC | 7 | SRSF5,SNRNP70,PABPN1,CCNL2,DDX39B,ITPKC,ARGLU1 | $4.807\times10^{-8}$ |
| Positive regulation of transcription by RNA | GO:0045944 | BP | 8 | NME2,DCN,PER1,EHF,EEF1D,ABLIM1,LPIN1,RARG | $2.111\times10^{-5}$ |
| Signal transduction | GO:0007165 | BP | 10 | TRIP10,ANXA1,ECM1,STK24,IMPA2,DUSP1,SFN,ARHGAP27,MAP3K6,RAPGEFL1 | $4.155\times10^{-3}$ |
| Intracellular signal transduction | GO:0035556 | BP | 6 | STK24,WSB1,DUSP1,HSPB1,DGKA,MKNK2 | $6.990\times10^{-3}$ |
| RNA processing | GO:0006396 | BP | 1 | PABPN1 | |
| RNA splicing | GO:0008380 | BP | 3 | ESRP2,DDX39B,ARGLU1 | |
| RNA catabolic process | GO:0006401 | BP | 1 | RNH1 | |
| Regulation of DNA-templated transcription | GO:0006355 | BP | 1 | BHLHE40 | |
| Transcription by RNA polymerase II | GO:0006366 | BP | 1 | ABLIM1 | |
| Chromatin remodeling | GO:0006338 | BP | 1 | PER1 | |
| Cytoplasmic stress granules | G0: 0010494 | CC | 1 | CIRBP | |
| PML body | GO:0016605 | CC | 1 | MKNK2 | |
| Centrosome | GO:0005813 | CC | 3 | EPS8L2,IL1RN,CALML3 | |
| Cytoplasmic ribonucleoprotein | GO:0036464 | CC | 1 | PABPN1 | |

**B**

| GO Term | GO ID | Category | Number of genes | Representative genes | p-value adjusted |
|---|---|---|---|---|---|
| Nuclear speck | GO:0016607 | CC | 12 | PNISR, DDX39B, SNRNP70, CCNL2, SRSF5, SRSF11, NXF1, SGK1, ATF4, PABPN1, NR4A1, ARGLU1 | $3.75\times10^{-14}$ |
| Positive regulation of transcription by RNA polymerase II | GO:0045944 | BP | 10 | PKD1, ATF4, JUN, CDK5RAP3, PLPP3, EGR1, CCN1, NR4A1, EEF1D, CALCOCO1 | $1.04\times10^{-07}$ |
| Cytoplasmic stress granules | G0: 0010494 | CC | 3 | NXF1, RBPMS | $1.32\times10^{-05}$ |
| Signal transduction | GO:0007165 | BP | 10 | PKD1, PLPP3, RHOB, SPARCL1,PPP1R12C, DUSP1, CCN1, NR4A1, CALCOCO1, IGFBP5 | $4.15\times10^{-03}$ |
| Centrosome | GO:0005813 | CC | 3 | SORBS1, ATF4, CDK5RAP3 | $7.03\times10^{-03}$ |
| Regulation of DNA-templated transcription | GO:0006355 | BP | 1 | ATF4 | |
| P-body | GO:0000932 | CC | 1 | RBPMS | |
| Transcription by RNA polymerase II | GO:0006366 | BP | 2 | NR4A1, ATF4 | |
| RNA splicing | GO:0008380 | BP | 4 | ARGLU1, DDX39B, SRSF11, RBM5 | |
| Intracellular signal transduction | GO:0035556 | BP | 7 | DUSP1, WSB1, SOCS3, IGFBP5, PKD1, SGK1 | |
| RNA processing | GO:0006396 | BP | 3 | RMB5, PABPN1, RBPMS | |

*Supplementary table 2. Gene Ontology (GO) enrichment analysis of downregulated genes in (A) Esophagous and (B) gastric tumors*

### ESOPHAGUS 2FOLD

#### 1. Proteins with Csat < 10μM: 28 proteins

- MED12: $C_{sat}$ = 0.07μM, Disordered residues = 1034
- ASCC3: $C_{sat}$ = 0.10μM, Disordered residues = 429
- CHERP: $C_{sat}$= 0.18μM, Disordered residues = 496
- MED13: $C_{sat}$= 0.19μM, Disordered residues = 1292
- PHIP: $C_{sat}$= 0.21μM, Disordered residues = 759
- LMBR1: $C_{sat}$= 0.42μM, Disordered residues = 104
- NEDD4L: $C_{sat}$= 0.58μM, Disordered residues = 368
- SP100: $C_{sat}$ = 0.60μM, Disordered residues = 571
- ZC3H13: $C_{sat}$ = 0.76μM, Disordered residues = 1341
- SCAF11: $C_{sat}$ = 0.78μM, Disordered residues = 1323
- MAP4K4: $C_{sat}$ = 0.85μM, Disordered residues = 517
- RIF1: $C_{sat}$ = 1.06μM, Disordered residues = 1507
- CDK12: $C_{sat}$ = 1.22μM, Disordered residues = 1145
- LATS1: $C_{sat}$ = 1.44μM, Disordered residues = 749
- ZBTB38: $C_{sat}$ = 2.33μM, Disordered residues = 1003
- NCOR2: $C_{sat}$ = 2.55μM, Disordered residues = 2214
- IRS1: $C_{sat}$ = 2.62μM, Disordered residues = 1031
- ARHGAP21: $C_{sat}$ = 2.77μM, Disordered residues = 1579
- NCOR1: $C_{sat}$ = 3.33μM, Disordered residues = 2138
- ICE1: $C_{sat}$ = 3.42μM, Disordered residues = 1747
- RTKN: $C_{sat}$ = 3.45μM, Disordered residues = 251
- GRB10: $C_{sat}$ = 4.14μM, Disordered residues = 215
- TNRC6B: $C_{sat}$ = 4.22μM, Disordered residues = 1705
- ATRX: $C_{sat}$ = 4.29μM, Disordered residues = 1649
- MAP3K2: $C_{sat}$ = 4.78μM, Disordered residues = 276
- TOPBP1: $C_{sat}$ = 5.93μM, Disordered residues = 680
- ETV6: $C_{sat}$ = 6.11μM, Disordered residues = 275
- ABL2: $C_{sat}$ = 8.77μM, Disordered residues = 653

#### 2. Proteins with Csat 10-50μM AND >500 disordered residues: 8 proteins

- MED1: $C_{sat}$ = 10.83μM, Disordered residues = 1149
- RAPGEF1: $C_{sat}$ = 12.58μM, Disordered residues = 576
- HCFC1: $C_{sat}$ = 17.49μM, Disordered residues = 1513
- BMPR2: $C_{sat}$ = 18.37μM, Disordered residues = 644
- COL15A1: $C_{sat}$ = 21.07μM, Disordered residues = 973
- MAGED1: $C_{sat}$ = 26.07μM, Disordered residues = 558
- USF3: $C_{sat}$ = 26.50μM, Disordered residues = 2144
- MAP1S: $C_{sat}$ = 38.98μM, Disordered residues = 519

### ESOPHAGUS 4FOLD

#### 1. Proteins with Csat < 10μM: 2 proteins

- AHR: $C_{sat}$ = 0.80μM, Disordered residues = 558
- IRS2: $C_{sat}$ = 7.20μM, Disordered residues = 1132

| 2. Proteins with Csat 10-50μM AND >500 disordered residues: 4 proteins |
|---|
| - FOXM1: $C_{sat}$ = 13.43μM, Disordered residues = 619 |
| - PPFIA1: $C_{sat}$ = 14.14μM, Disordered residues = 510 |
| - WNK2: $C_{sat}$ = 28.36μM, Disordered residues = 1860 |
| - MYBL2: $C_{sat}$ = 34.10μM, Disordered residues = 504 |

*Supplementary table 4. Results from the ML predictor indicating the obtained $C_{sat}$ values and the number of disordered amino acids in esophageal cancer. The table is divided in 2-fold and 4-fold, separating those sequences with $C_{sat}$<10μM and $C_{sat} \in (10,50)$ μM with more than 500 disordered residues.*
